# Phosphorylation of phosphoglucomutase 1 on a peripheral site tunes its activity to regulate glycogen metabolism

**DOI:** 10.1101/2021.04.15.439997

**Authors:** Sofía Doello, Niels Neumann, Karl Forchhammer

**Affiliations:** Interfaculty Institute of Microbiology and Infection Medicine, University of Tübingen, Auf der Morgenstelle 28, 72076 Tübingen, Germany

**Keywords:** glycogen metabolism, phosphoglucomutase, phosphorylation, cyanobacteria

## Abstract

Regulation of glycogen metabolism is of vital importance in organisms of all three kingdoms of life. Although the pathways involved in glycogen synthesis and degradation are well known, many regulatory aspects around the metabolism of this polysaccharide remain undeciphered. Here, we used the unicellular cyanobacterium *Synechocystis* as a model to investigate how glycogen metabolism is regulated in nitrogen-starved dormant cells, which entirely rely on glycogen catabolism to resume growth upon nitrogen repletion. We identified phosphoglucomutase 1 (PGM1) as a key regulatory point in glycogen metabolism, and post-translational modification as an essential mechanism for controlling its activity. We could show that PGM1 is phosphorylated at a peripheral residue (Ser 47) during nitrogen starvation, which inhibits its activity. Inactivation of PGM1 by phosphorylation at Ser 47 prevents premature degradation of the glycogen stores and appears to be essential for survival of *Synechocystis* in the dormant state. Remarkably, this regulatory mechanism seems to be evolutionary conserved in PGM1 enzymes, from bacteria to humans.

**Significance statement:** In this study, we identified phosphoglucomutase 1 (PGM1) as a central metabolic valve that regulates the utilization of the glycogen reserves. We showed that post-translational modification of PGM1 via phosphorylation at a peripheral residue is a key, evolutionary-conserved regulatory mechanism that controls PGM1 activity and the mobilization of the glycogen stores.

## Introduction

Glycogen is the major carbohydrate storage compound in a broad range of organisms, from bacteria to humans. This polysaccharide is composed of glucose molecules connected by α,1-4 linkages and branched via α,1-6 linkages, and it is generally considered a carbon sink with energy-storage function. In humans, glycogen is mainly accumulated in the liver and skeletal muscle, and it constitutes a rapid and accessible form of energy that can be supplied to tissues on demand.^1^ In many bacteria, glycogen plays a crucial role in survival to an ever-changing environment. It is usually synthesized and accumulated inside the cells under growth-limiting conditions at excess of a carbon source, and degraded when the supply of energy or carbon is not enough to maintain growth or viability, thus allowing cell survival in transient starvation conditions.^2^ In cyanobacteria, which generally sustain cell growth by performing oxygenic photosynthesis, glycogen is synthesized towards the end of the day, when photosynthetically fixed carbon is in excess and cells need to prepare to survive the night.^3^ Glycogen accumulation also occurs as a response to nutrient limitation. In fact, the greatest amount of glycogen accumulation in non-diazotrophic cyanobacteria, which are unable to fix atmospheric N2, occurs under nitrogen starvation conditions.^4^

Nitrogen deprivation activates a genetically determined survival program in non-diazotrophic cyanobacteria, which has been extensively studied in the unicellular cyanobacterial strains *Synechococcus elongatus* and *Synechocystis* sp. PCC 6803 (from now *Synechocystis*).^5,6^ When *Synechocystis* encounters nitrogen depletion, the intracellular carbon/nitrogen balance is disturbed, and growth can no longer be supported. This metabolic situation leads to rapid accumulation of glycogen, which serves as a sink for the excess of carbon.^5^ In order to survive these starvation conditions, cells undergo an adaptation process termed chlorosis that involves the degradation of the light-harvesting complexes to avoid an excess of energy and reduction equivalents that are no longer consumed by anabolic reactions. As a result of the metabolic and morphological changes induced by nitrogen starvation, cells enter a dormant state, which allows them to survive adverse conditions for a prolonged period of time.^6^ Upon nitrogen availability, the glycogen stores accumulated in dormant cells play a key role in the restoration of vegetative growth.^7^ When dormant cells have access to a nitrogen source, their metabolism switches towards a glycolytic phase. They turn off residual photosynthesis, while the production of energy and metabolic intermediates now entirely relies on glycogen catabolism.^8^ This extraordinary situation, in which carbohydrate degradation can be completely separated from photosynthetic processes even in the presence of light, makes awakening *Synechocystis* cells an excellent model to study the regulation of glycogen catabolism.

Although the metabolic pathways involved in glycogen synthesis and degradation are well known, many regulatory aspects around the metabolism of this polysaccharide remain to be deciphered. In nitrogen-starved *Synechocystis* cells, glycogen degradation is known to start soon after addition of a nitrogen source, and the enzymes responsible for this process have been identified (**Figure 1**).^7^ However, how glycogen catabolism is induced in dormant cells has not yet been elucidated. The enzymes involved in glycogen metabolism are conserved from bacteria to humans. The glycogen phosphorylase and debranching enzyme are responsible for the excision of glucose molecules from the glycogen granule, releasing glucose-1-phosphate (glucose-1P) and glucose, respectively. Glucose-1P is then converted to glucose-6-phosphate (glucose-6P) by the phosphoglucomutase (PGM), an evolutionary conserved enzyme that also catalyzes the reverse reaction, while glucose is converted to glucose-6P by the glucokinase. Glucose-6P can then enter different catabolic pathways to produce glyceraldehyde-3-phosphate. Intriguingly, most of the glycogen catabolic enzymes are up-regulated during nitrogen starvation, although glycogen degradation does not start until a nitrogen source is available. This suggests that the activity of these enzymes must be tightly regulated: They must remain inactive when cells are dormant and be activated upon nitrogen availability. An exception to the abundance pattern of most glycogen catabolic enzymes is PGM1, the expression of which is suppressed under nitrogen starvation and activated during resuscitation.^8,9^ Although *Synechocystis* possesses two PGM isoenzymes, PGM1 (*sll0726*) has been shown to be responsible for almost 97 % of the PGM activity,^10^ while PGM2 (*slr1334*) serves as glucose-1,6-bisphosphate synthase.^11^ PGM1 was recently identified as a phosphoprotein with two localized serine phosphorylation sites: Ser 47 and Ser 152. Ser 152 corresponds to the catalytically active serine residue, which mediates the phosphoryl transfer between the sugar and the enzyme. In typical PGM enzymes, the catalytic serine residue is phosphorylated by the activator glucose-1,6-bisphosphate.^11^ During chlorosis, the degree of phosphorylation of this residue decreases. On the contrary, the phosphorylation of Ser 47 strongly increases during nitrogen starvation, representing one of the most strongly induced phosphorylation events.^9^ The function of this serine residue is so far unknown; previous studies have not considered it to be involved in catalysis or regulation of PGM1 activity.^12,13^ These findings prompted us to investigate the possible involvement of PGM1 in the regulation of glycogen metabolism, leading to the identification of a key regulatory mechanism in glycogen catabolism, which seems to be conserved from bacteria to humans.

**Figure 1.**
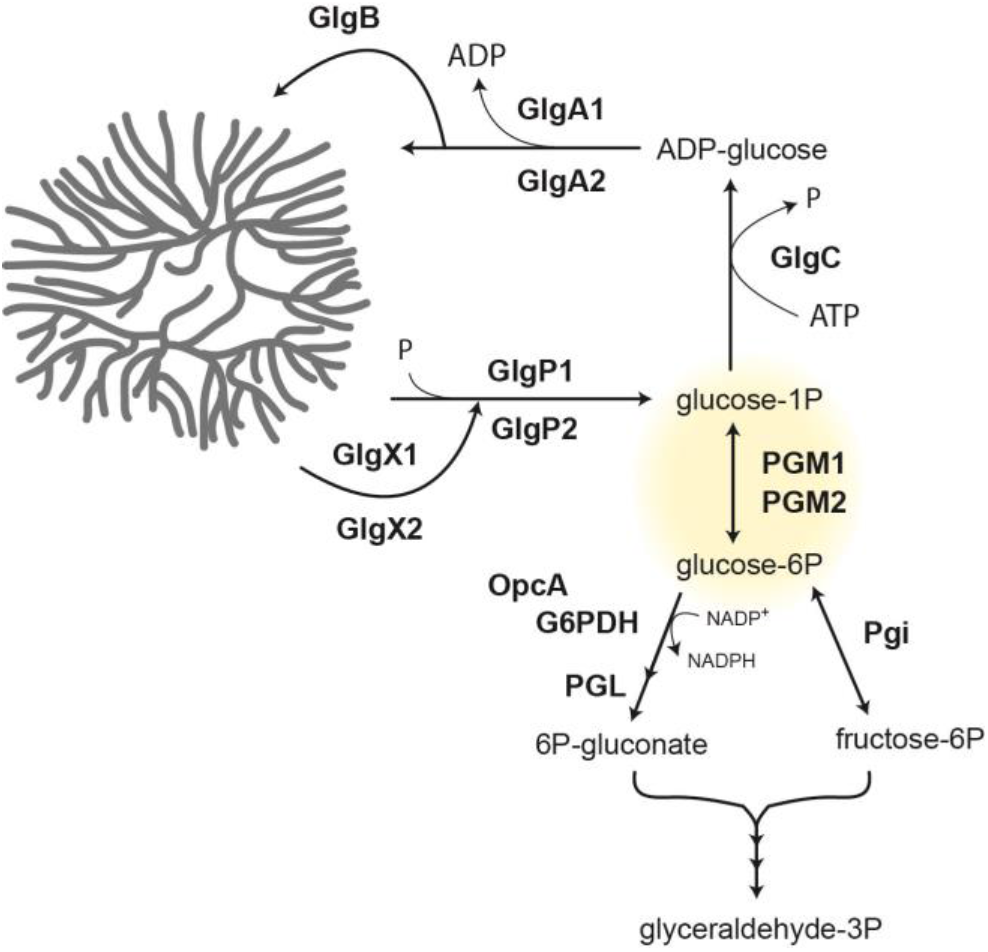
Schematic representation of the glycogen metabolic hub in *Synechocystis*. Glycogen is synthesized by the glycogen synthases (GlgAs) and branching enzyme (GlgB) from ADP-glucose, which is obtained from glucose-1-phosphate (glucose-1P) through the reaction catalyzed by the glucose-1-phosphate adenylyltransferase (GlgC). Glycogen catabolism starts with the glycogen phosphorylases (GlgPs) and debranching enzymes (GlgXs). The glucose-1P released by GlgPs is converted to glucose-6-phosphate (glucose-6P) by the phosphoglucomutases (PGMs). Glucose-6P can be metabolized by the phosphoglucoisomerase (Pgi) to enter the Embden-Meyerhof-Parnas pathway or by the glucose-6P dehydrgogenase (G6PDH), which requires the activator protein OpcA, to enter the oxidative pentose phosphate pathway and the Enter-Doudoroff pathway. Ultimately, all pathways generate glyceraldehyde-3-phosphate (glyceraldehyde-3P).

## Results

### PGM1 is activated during resuscitation from nitrogen starvation

Our previous studies suggested the occurrence of an unknown mechanism controlling the activity of glycogen catabolic enzymes in *Synechocystis*: The transcription of most glycogen catabolic genes is highly up-regulated during nitrogen deprivation when glycogen is synthesized, but turned down during resuscitation when glycogen is catabolized.^14,7^ In agreement, a proteomic study showed increased abundance of these enzymes in the dormant state and return to normal levels during resuscitation (**Figure 2A**).^9^ One exception to this expression pattern is PGM1, the abundance of which is low during nitrogen starvation and increases during resuscitation (**Figure 2A**). In the same study, a quantitative analysis of the phosphorylation events during nitrogen starvation and resuscitation revealed that PGM1 can be phosphorylated at two different serine residues. Due to an incorrect annotation of PGM1 in CyanoBase,^15^ the residues previously designated as Ser 63 and Ser 168^9^ correspond to the residues Ser 47 and Ser 152 of PGM1 (see below, **Figure S2**). Interestingly, Ser 47 is one of the most strongly phosphorylated residues in chlorotic cells, being 13 times more phosphorylated under nitrogen starvation than during vegetative growth (**Figure 2B**). These findings suggested that PGM1 might be a regulatory point, ensuring that glycogen catabolism is arrested in the chlorotic state until nitrogen becomes available again.

**Figure 2.**
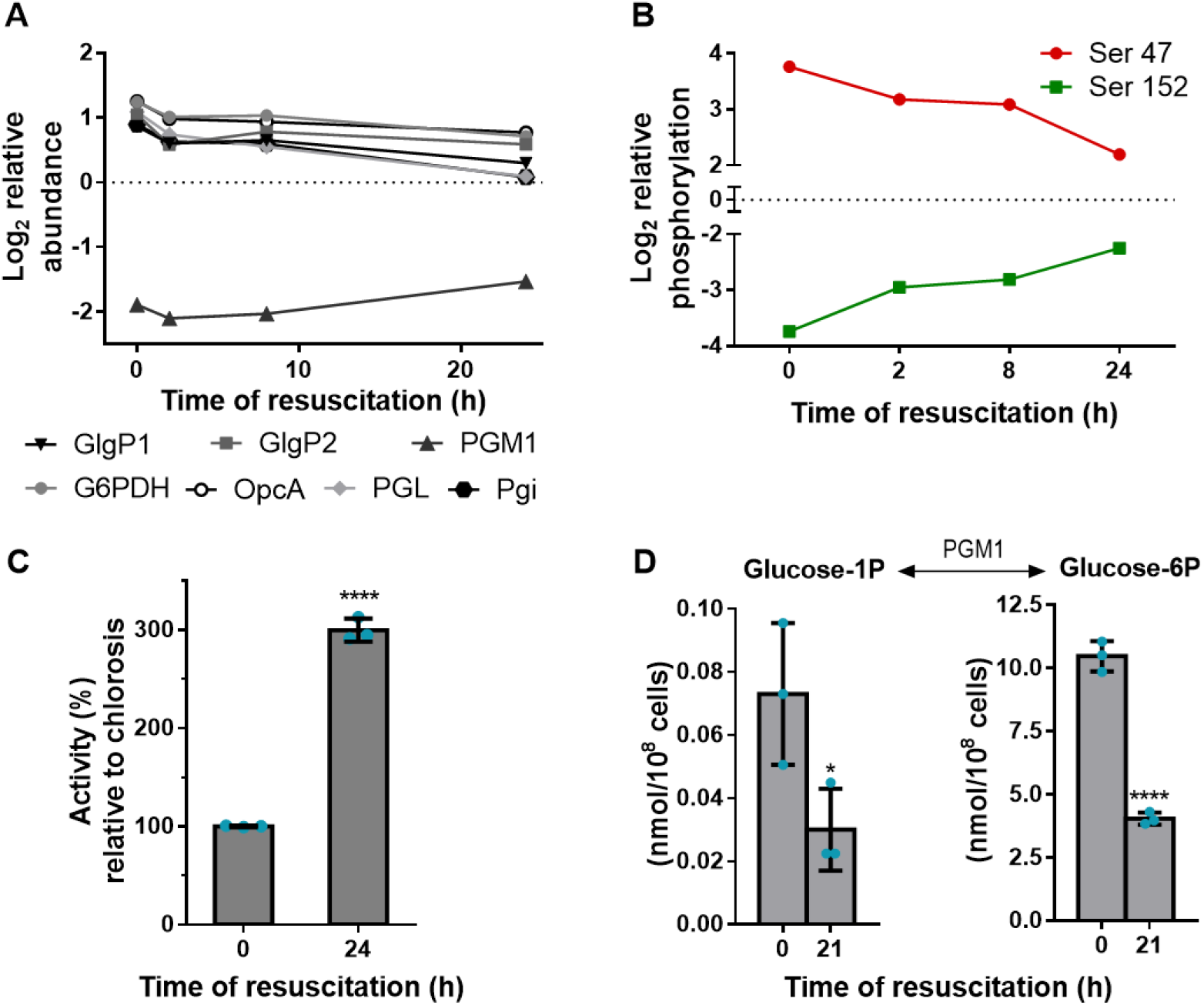
Abundance, phosphorylation and activity of PGM1 during nitrogen starvation and resuscitation. (A) Protein abundance ratios of GlgP1, GlgP2, PGM1, G6PDH, OpcA, PGL, and Pgi during resuscitation from nitrogen starvation. Ratios were calculated comparing the protein abundance during nitrogen starvation and resuscitation with their abundance during vegetative growth. Relative abundance is shown as the log_2_ of the calculated ratios. Positive values indicate up-regulation and negative values down-regulation compared to protein levels during vegetative growth (normalized to zero, dotted line).^9^ (B) Phosphorylation events of the two phosphorylation sites in PGM1 at the indicated time points during resuscitation from nitrogen starvation. Ratios were calculated comparing the abundance of phosphorylated and unphosphorylated peptides at different time points during resuscitation to their abundance during vegetative growth. Relative phosphorylation is shown as the log_2_ of the calculated ratios.^9^ (C) Relative enzyme activity of PGM1 in cell extracts from chlorotic and resuscitating cells. The activity in chlorotic cells was considered as 100%. (D) Glucose-1P and glucose-6P content normalized to 10^8^ cells during nitrogen starvation and resuscitation. For (C) and (D) at least three biological replicates were measured. Error bars represent the standard deviation (SD); asterisks represent the statistical significance.

To test whether there was any change in the activity of PGM1 upon addition of a nitrogen source to dormant cells, we assayed PGM1 activity in cell extracts from chlorotic and resuscitating cells (**Figure 2C**). While some PGM1 activity was detectable in nitrogen-starved cells, the measured activity was three times higher in cells supplemented with nitrate 24 hours before. These results suggested an activation of PGM1 upon addition of nitrogen to chlorotic cells. Analysis of the levels of glucose-phosphates during nitrogen starvation and resuscitation supported this idea because higher levels of glucose-1P and glucose-6P, the substrates of PGM1, were detected in chlorotic than in recovering cells (**Figure 2D**). Given the significant phosphorylation changes detected in PGM1 during nitrogen starvation, we hypothesized this post-translational modification could regulate enzyme activity in the different developmental stages.

### PGM1 activity is regulated via phosphorylation at Ser 47

According to homology modeling of PGM1, Ser 152 is the catalytic serine residue involved in the phosphor-exchange reaction. This residue is highly conserved in the members of the α-D-phosphohexomutase (α-D-PHM) superfamily, and it is located in the active site of PGM1 (marked in green in **Figure 3A**). This catalytic serine is poorly phosphorylated during nitrogen starvation, and it progressively becomes more phosphorylated during resuscitation (**Figure 2B**). Since phosphorylation of the catalytic serine is required for catalysis, the phosphorylation dynamics of this residue corresponds to its state of catalytic activity, with PGM1 being inactive in chlorotic cells and becoming activated during resuscitation. Ser 47 follows the opposite pattern: The high level of phosphorylation of this residue under nitrogen starvation progressively decreases during resuscitation (**Figure 2B**). As deduced from homology modeling, Ser 47 is located on the surface of the enzyme, in a loop known as “the latch” (marked in red in **Figure 3A**). The phosphorylation dynamics of Ser 47 was confirmed using clear-native PAGE – immunoblot analysis. Because Ser 47 is on the surface of the enzyme, addition of a phosphate group to this residue affects the net charge of the protein and influences its migration in a native gel, whereas phosphorylation of Ser 152, which is buried in the catalytic center, does not. As shown in **Figure 3B,** after 24 h of nitrogen starvation a lower band appeared in the gel, which represents PGM1 phosphorylated at Ser 47 (PGM1^2^). After 7 days of chlorosis, only the lower band is observed. The upper band, which represents PGM1 dephosphorylated at Ser 47 (PGM1^1^), is only observed again 8 h after resuscitation was initiated. To confirm that the different bands correspond to different phosphorylation states of PGM1, cell extracts from chlorotic cells were prepared in the absence of phosphatase inhibitors and treated with alkaline phosphatase. Under these conditions, most of PGM1 was detected in the upper band, corresponding to the non-phosphorylated state (**Figure 3B**).

**Figure 3.**
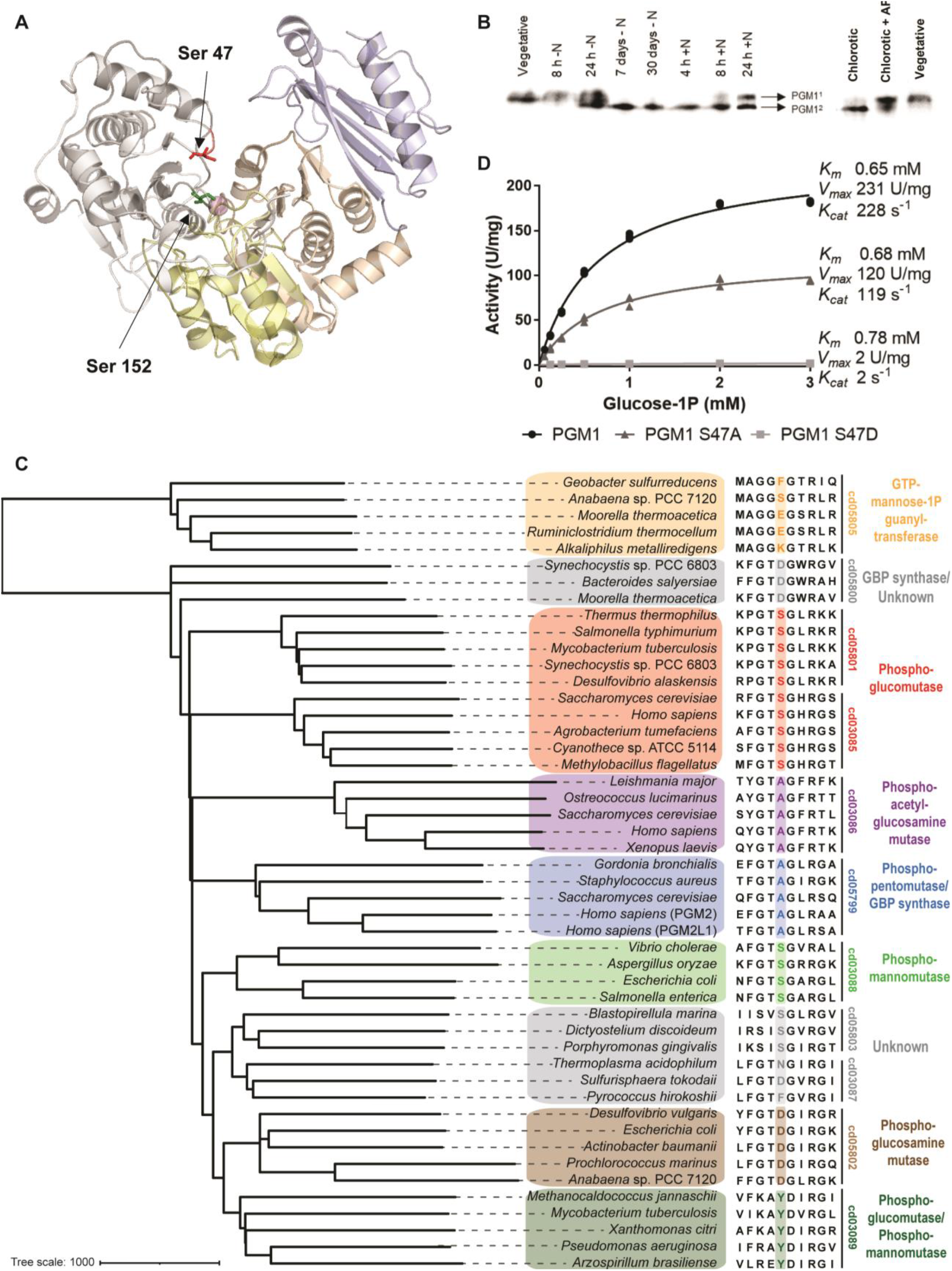
Phosphorylation of Ser 47 regulates PGM1 activity. (A) Structure of *Synechocystis’* PGM1 obtained from Swiss Model using *Salmonella typhimurium’*s PGM1 as a template (Protein Data Bank code 3OLP). The two colored resides shown in a stick model represent the two phosphorylation sites: Ser 47 (and the latch domain) in red and Ser 152 in green. The Mg^+^ ion required for catalysis is shown as a pink sphere. Domain 1 is colored in grey, domain 1 in yellow, domain 3 in orange, and domain 4 in blue. (B) Clear native PAGE - immunoblot validation of PGM1 phosphorylation dynamics during nitrogen starvation, resuscitation, and after treatment with 1 U/mL of alkaline phosphatase (AP) for 10 min. PGM1^1^ and PGM1^2^ represent PGM1 dephosphorylated and phosphorylated at Ser 47, respectively. (C) Phylogenetic tree of the α-D-PHM superfamily showing conservation of the residue Ser 47 of *Synechocystis*. (D) Michaelis-Menten kinetics of wild type (WT) PGM1 (black circles), PGM1 S47A (dark grey triangles), and PGM1 S47D (light grey squares). Three replicates were measured for each data-point. See also Figures S1 and S2.

While the residue Ser 152 in *Synechocystis* PGM1 is conserved throughout the α-D-PHM superfamily, since it directly participates in catalysis, Ser 47 appears to be subfamily-specific. The α-D-PHM superfamily comprises different classes of enzymes, which share a common structure and mechanism of catalysis.^16^ The Conserved Domain Database (CDD) classifies the members of the α-D-PHM superfamily into different subfamilies according to their domain architecture.^17^ Based on their substrate specificities, these subfamilies can be grouped into: phosphoglucomutases or PGM1 (cd03085 and cd05801), phosphomannomutases or PMM (cd03088), bi-functional phosphogluco/phosphomanno mutases or PGM/PMM (cd03089), phosphoglucosamine mutases or PNGM (cd05802), phosphoacetylglucosamine mutases or PAGM (cd03086), phosphopentomutases/glucose-1,6-bisphosphate synthases or PGM2/PGM2L1 (cd05799), and enzymes of unknown function (cd03087, cd05800, and cd05803). The residue Ser 47 is located in the latch loop and as shown in the phylogenetic tree in **Figure 3C**, it is conserved in the members of the subfamilies cd05801 and cd03085, which are PGM1 enzymes. According to structural studies on enzymes from the cd05801 and cd03089 subfamilies, this loop participates in a conformational change required for catalysis and in stabilization of the substrate through interaction with the phosphorylated end of the sugar.^18–21^ The presence of a residue with a phosphorylatable hydroxyl group in its side chain at the position of Ser 47 might be a determinant for phosphoglucomutase reaction velocity since the subfamilies that have high phosphoglucomutase activity (cd03085, cd05801, cd03088 and cd03089)^13,19,21–24^ present Ser or Tyr at this site, while the subfamilies presenting Ala (cd03086, and cd05799) or Asp (cd05802) at this position possess low phosphoglucomutase activity (**Figure S1**).^11,25–27^

To gain more insights on the role of Ser 47 in catalysis, we created different recombinant PGM1 variants with site-specific amino acid substitutions. Analysis of the sequences of PGM1 from several *Synechocystis* strains revealed a different annotation for *Synechocystis* sp. PCC 6803, which included a 16-amino acid N-terminal extension that is missing in the other strains (**Figure S2A**). Additionally, the experimentally validated transcriptional start site from *Synechocystis* sp. PCC 6803 suggests a shorter open reading frame for PGM1, with Met 17 as putative translational start site.^28^ To clarify if the N-terminal extension is an annotation error, we prepared as recombinant proteins the PGM1 as annotated in Cyanobase^15^ (Long-PGM1) and a version without the 16-amino acid N-terminal extension (Short-PGM1). We then measured their activity *in vitro*, assuming that the native version exhibits higher activity. Although both versions showed a similar substrate affinity, as deduced from the calculated Michaelis-Menten constant (*K_m_*), the maximal velocity (*V_max_*) of the reaction was almost 3-fold higher for the short version (**Figure S2B**), substantiating that the short version likely represents the physiologically relevant protein. The different PGM1 variants were therefore created from the Short-PGM1 version (hereafter referred to as PGM1). To elucidate the importance of the interactions of the hydroxyl group of Ser 47 in catalysis, this residue was replaced by Ala (PGM1 S47A), a smaller and hydrophobic residue. Comparison of the kinetic parameters of the wild-type (WT) PGM1 and PGM1 S47A showed that the substitution of Ser for Ala at position 47 affected the *V_max_* of the reaction, which decreased almost 2-fold, but it did not decrease substrate affinity, as shown by the *K_m_* value (**Figure 3C**). These results indicate that the interactions in which Ser 47 is involved are not essential but play a modulating role in catalysis. Phosphorylation of this residue could therefore disturb these interactions and inhibit PGM1 activity. To estimate the effect of phosphorylation of this residue on enzyme activity, Ser 47 was substituted by Asp (PGM1 S47D), a larger and negatively charged amino acid that resembles a phosphorylated Ser. PGM1 S47D presented very low activity *in vitro* (0.86 % of the WT activity, **Figure 3C**), showing that phosphorylation of Ser 47 inactivates PGM1.

### Phosphorylation of PGM1 at Ser 47 is essential for survival under nitrogen starvation

To determine the physiological significance of phosphorylation of Ser 47 during nitrogen starvation in *Synechocystis*, we created and characterized various PGM1 mutant strains. A PGM1 knockout strain (Δ*pgm1*) could not properly acclimate to nitrogen-depletion and presented a so-called non-bleaching phenotype: Cells did not degrade their photosynthetic pigments and turned yellow, but stayed greenish instead, progressively looking paler (**Figure S3A**). After seven days of nitrogen starvation, a very reduced proportion of cells could recover when they were dropped on an agar plate containing nitrate, as compared to the WT (**Figure S3B**). Such a phenotype was previously observed in mutants that were impaired in glycogen synthesis,^29^ since accumulation of this polymer has been shown to be indispensable for adaptation to nitrogen-starvation. This phenotype was expected, given that PGM1 catalyzes the interconversion between glucose-1P and glucose-6P and is therefore involved in glycogen synthesis. Indeed, no glycogen was detected in seven-days-starved Δ*pgm1* cells (**Figure S3C**), indicating that PGM1 activity is essential for glycogen synthesis under nitrogen deprivation and that the activity of PGM2 does not compensate the lack of PGM1. This is consistent with the fact that phosphorylation of Ser 47 is only detected after 24 h of nitrogen depletion (**Figure 3B**), when synthesis of the glycogen stores has already taken place. Consequently, a strain with an inactive PGM1 variant, such as the PGM1 S47D, would not be able to enter the chlorotic state due to its inability to synthesize glycogen. To study the physiological consequences of the lack of PGM1 inactivation via phosphorylation, we complemented the Δ*pgm1* strain with the WT PGM1 (Δ*pgm1*+PGM1) and with the partially active PGM1 S47A variant (Δ*pgm1*+PGM1S47A), which lacks the phosphorylation site. Complementation with the WT protein rescued the phenotype: The Δ*pgm1*+PGM1 strain showed a similar behavior than the WT under nitrogen starvation (**Figure S4**). Conversely, the Δ*pgm1*+PGM1S47A strain could not successfully enter chlorosis. Although it seemed to induce pigment degradation upon nitrogen starvation (in contrast to the Δ*pgm1* strain), these cultures did not acquire the same characteristic yellowish color than the WT, but a pale color instead (**Figure 4A**) and showed very poor recovery on nitrate-containing agar plates (**Figure 4B and C**). This strain could not accumulate the same amount of glycogen than the WT (**Figure 4D**), presumably due to the lower activity of the PGM1 S47A variant, which was not enough to meet the cellular demand. Since the Δ*pgm1*+PGM1S47A strain failed to properly acclimate to nitrogen deprivation, we transformed WT cells with PGM1 S47A (WT+PGM1S47A) to study the impact of the presence of a PGM1 variant lacking phosphorylation of residue 47 on long-term nitrogen starvation. As expected, this strain could initially acclimate to nitrogen depletion like the WT (**Figure 4A**), since it contains the WT version of PGM1. After 6 days of chlorosis, WT+PGM1S47A cells could recover as efficiently as WT cells on nitrate-containing agar plates (**Figure 4B**). However, after prolonged exposure to nitrogen deprivation, in which the WT version of PGM1 would be highly phosphorylated, the cultures of the WT+PGM1S47A strain progressively lost their yellowish color (**Figure 4A**). After one month of starvation, only a reduced number of cells could recover upon nitrogen repletion (**Figure 4C**). WT+PGM1S47A cells could synthesize a similar amount of glycogen than WT cells upon nitrogen depletion, but after one week of starvation the glycogen content began to gradually decrease (**Figure 4D**). These findings imply that inactivation of PGM1 via phosphorylation is crucial for preventing premature glycogen degradation during prolonged nitrogen starvation, which appears to be essential for survival of these conditions.

**Figure 4.**
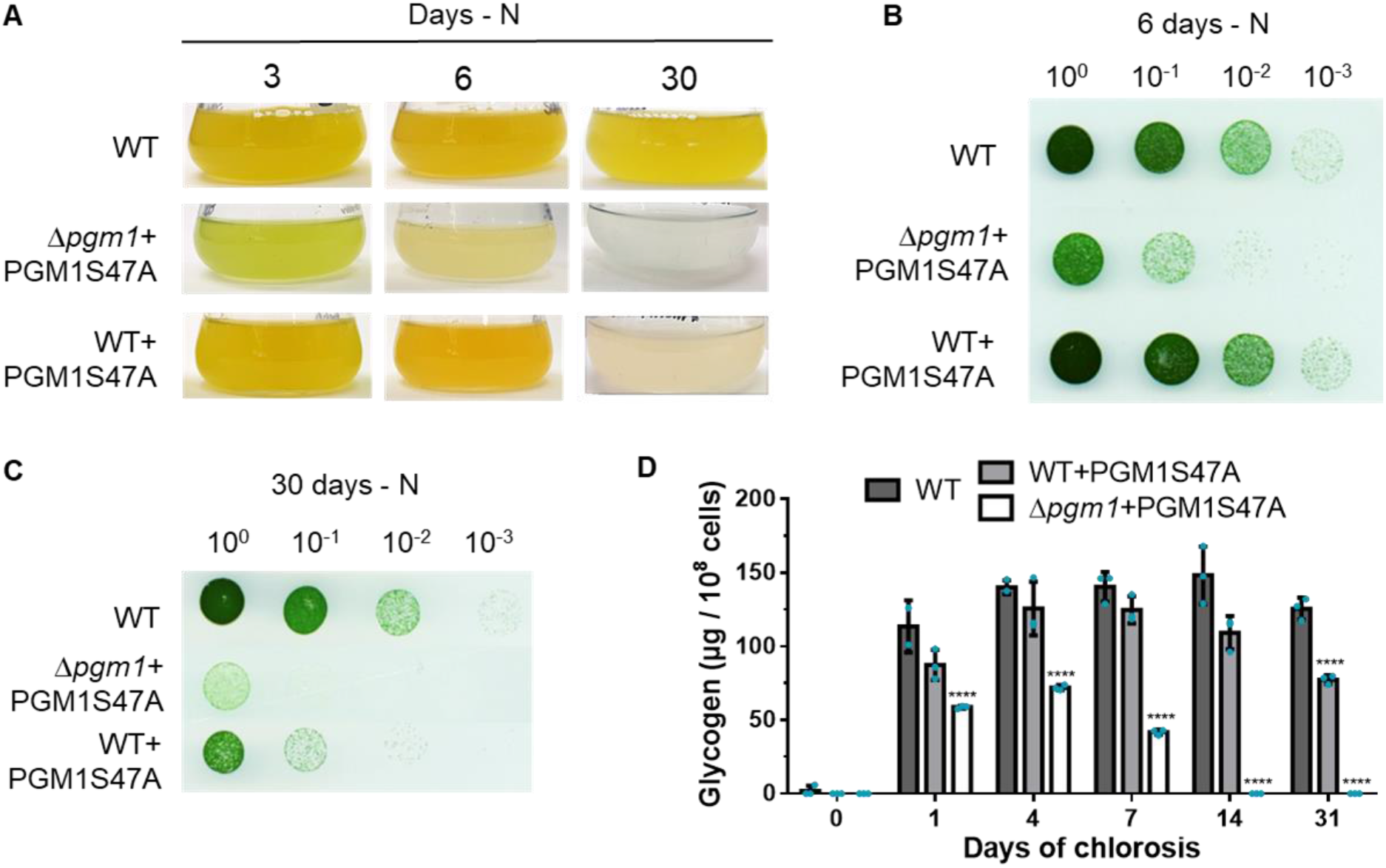
PGM1 is required for glycogen synthesis, and glycogen degradation during nitrogen starvation is prevented by phosphorylation of Ser 47. (A) Pictures of WT, Δ*pgm1*+PGM1S47A, and WT+PGM1S47A cultures after 3, 6, and 30 days of nitrogen starvation. (B) Recovery assay on a BG_11_-agar plate of WT, Δ*pgm1*+PGM1S47A, and WT+PGM1S47A after 6 days of nitrogen starvation. Numbers on top represent the dilution factor, starting with an OD_750_ of 1. Pictures were taken 5 days after dropping chlorotic cells on the plate. (C) Recovery assay on a BG_11_-agar plate of WT, Δ*pgm1*+PGM1S47A, and WT+PGM1S47A after 30 days of nitrogen starvation. Numbers on top represent the dilution factor, starting with an OD_750_ of 1. Pictures were taken 5 days after dropping chlorotic cells on the plate. (D) Glycogen content of WT, Δ*pgm1*+PGM1S47A and WT+PGM1S47A at the indicated time points during nitrogen starvation. In all experiments, three biological replicates were measured. Error bars represent the SD; asterisks represent the statistical significance. See also Figures S3 and S4.

### PGM1 regulation seems to be conserved from bacteria to humans

Regulation of PGM1 activity is crucial for the survival of a wide range of organisms to many different conditions. Since all PGM1 enzymes present a phosphorylatable Ser residue at the site of Ser 47 of *Synechocystis*, regulation of PGM1 activity via phosphorylation of this residue might not be exclusive to *Synechocystis*, but widespread among the cd05801 and cd03085 subfamilies. Indeed, phosphorylation of this residue has been detected in the PGM1 from mammals (**Figure 5A**).^30,31^ In humans, PGM1 deficiency leads to glycogenosis, a metabolic disorder that causes the abnormal use and storage of glycogen, and congenital disorders of glycosylation. Despite the importance of the correct activity of this enzyme on human health, little is known regarding its regulation. Although Ser 20 of the human PGM1 (HPGM1), which is the homologous residue of *Synechocystis* PGM1 Ser 47 in the human enzyme, has been identified as a phosphorylation site,^30^ the role of phosphorylation of this residue has not been investigated. Lee et al.^32^ investigated the effects of mutations present in patients suffering from PGM1 deficiency on the *in vitro* activity of the PGM1 enzyme and identified six mutations that compromised catalysis and six mutations that caused folding defects (**Figure 5B**). Interestingly, a mutation in the latch loop (T19A) (**Figure 5C**) severely affected the catalytic activity of the enzyme,^32^ suggesting that addition of a phosphate group to an adjacent residue in this loop (i.e., S20) might also inhibit catalysis, as we observed for *Synechocystis* PGM1. To ascertain this, we prepared recombinant HPGM1 along with a mutant variant in which Ser 20 had been substituted by Asp (HPGM1 S20D) and measured their activities *in vitro*. As presented in **Figure 5D**, the phosphomimetic substitution of Ser 20 to Asp renders the enzyme inactive, indicating that the phosphorylated form of the enzyme is likely inactive.

**Figure 5.**
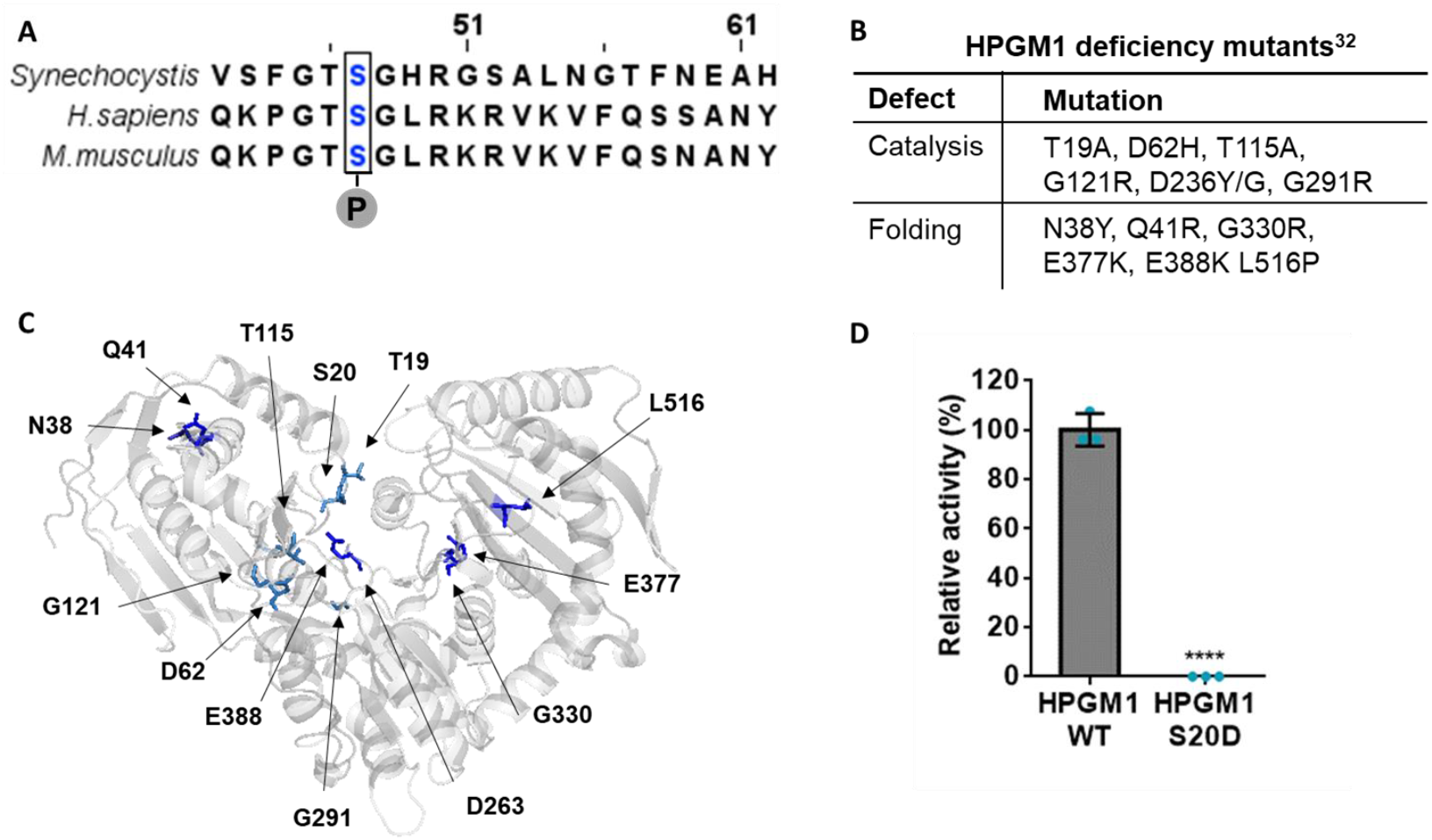
Regulation of PGM1 by phosphorylation is conserved in mammals. (A) Alignment of the sequence of PGM1 from *Synechocystis*, *Homo sapiens*, and *Mus musculus*. The conserved phosphorylation site is marked in blue. (B) List of human PGM1 (HPGM1) deficiency mutants and their associated defects.^32^ (C) Structure of HPGM1 (Protein Data Bank code 5EPC) with relevant residues highlighted. (D) Relative *in vitro* activity of HPGM1 and HPGM1 S20D. The activity of WT PGM1 was considered to be 100%. At least 3 replicates were measured. Error bars represent the SD; asterisks represent the statistical significance.

## Discussion

Regulation of glycogen metabolism is of vital importance in organisms of all three kingdoms of life. In bacteria, proper control of glycogen synthesis and degradation determines the ability to survive transient periods of nutrient starvation. In mammals, deficiencies in glycogen metabolism produce a variety of different metabolic disorders, some of them very severe.^32^ In this study, we used the unicellular cyanobacterium *Synechocystis* to investigate the regulation of glycogen metabolism. During nitrogen starvation, the activity of the glycogen catabolic enzymes must be regulated to prevent premature glycogen degradation, so that cells can use this reserve polymer once the conditions are favorable for resuscitation. We could show that phosphorylation of PGM1 at a peripheral residue regulates its activity and is required to prevent premature glycogen degradation.

PGM1 is an evolutionary conserved enzyme that mediates one of the most important reactions in carbohydrate metabolism: It catalyzes the interconversion between glucose-1P and glucose-6P, being thereby involved in both, glycogen synthesis and degradation.^33^ Despite its important role in sugar metabolism, PGM1 activity has only recently started to be considered as a target for metabolic control.^34–37^ At the onset of nitrogen starvation, the activity of PGM1 is required for glycogen synthesis, as shown by the inability of a PGM1 knock-out mutant to synthesize glycogen. Hence, during the first hours of nitrogen starvation, PGM1 remains dephosphorylated at Ser 47, a residue located in the so-called latch loop. Once enough glycogen has been accumulated, PGM1 activity is inhibited by phosphorylation of Ser 47. This residue is highly phosphorylated in long-term-chlorotic cells, and *in vitro* characterization of a phosphomimetic variant of PGM1 (PGM1 S47D) indicates that the phosphorylated version of the enzyme is inactive. When a non-phosphorylated PGM1 S47A variant is present in chlorotic cells, glycogen is degraded after prolonged nitrogen starvation, leading to loss of viability. This highlights the pivotal role of Ser 47 phosphorylation for controlling glycogen metabolism.

Analysis of the PGM1 S47A variant indicated that the interactions in which Ser 47 is involved are important for modulating catalysis, as the *V_max_* was 2-fold lower in the PGM1 S47A variant than in WT PGM1, without impairing the substrate affinity. This is in line with structural studies of the reaction mechanism on related proteins, which suggested a role of Ser 47 in a conformational change occurring during catalysis ^18–21^ When the enzyme is in its open conformation, its catalytic cleft is easily accessible for phosphorylated sugars. When the substrate enters the catalytic site, its dephosphorylated end interacts with the phosphorylated catalytic serine, and the phosphorylated end of the phosphor-sugar must interact with a group of residues from domain 4 (the phosphate-binding loop, colored in blue in **Figure 3A**). In order for these residues to come in contact with the substrate, the enzyme must undergo a conformational change that involves a rotation of domain 4 and changes the active site from an open cleft to a closed pocket. This conformational change requires the interaction of a group of residues from the phosphate-binding loop with the residues from the latch loop (colored in red in **Figure 3A**).^13,16,18,20,38^ When the hydroxyl group in the side chain of Ser 47 is eliminated by replacing it with Ala, the interactions that allow such conformational change and stabilization of the substrate are partially disturbed, which affects the velocity of the catabolic reaction. In the absence of the WT PGM1, a *Synechocystis* strain containing a non-phosphorylated PGM1 S47A variant accumulated only a reduced amount of glycogen, showing that the anabolic reaction is also affected with the substitution of Ser for Ala. When the site of Ser 47 is occupied by a negatively charged residue, as in the phosphomimetic variant or when the serine residue is phosphorylated, these interactions are completely disrupted, and catalysis is almost entirely inhibited.

Interestingly, Ser 47 is conserved among PGM1 enzymes (cd03085 and cd05801 subfamilies), and also in PMM enzymes (cd03088). In agreement with the finding that this residue is involved in modulating *K_cat_*, subfamilies that generally have high PGM1 activity ^13,19,21–24^ present a phosphorylatable amino acid (Ser or Tyr) at this site, while those subfamilies that show low PGM1 activity and preferably catalyze the interconversion of other phosphor-sugars/amino sugars (cd05802, cd03086, and cd05799) ^11,25–27^ contain Asp or Ala at this position. The Ser residue in the latch loop has also been identified as a phosphorylation site in the PGM1 from mammals, which belong to the cd03085 subfamily. Even though PGM1 deficiency can cause severe disease in humans, its regulation remains under-investigated. HPGM1 is known to be phosphorylated and thereby activated by the p21-activated signaling kinase 1 (Pak1) on Thr 466.^33^ However, although Ser 20, the homologous of *Synechocystis* Ser 47, had previously been reported as a phosphorylation site in HPGM1,^30^ the role of this phosphorylation on enzyme activity had not been characterized. We were able to demonstrate that, as in *Synechocystis*, HPGM1 is also inactivated by replacement of Ser 20 with a phosphomimetic residue. In agreement with our results, a study involving human patients suffering from PGM1 deficiency showed that introduction of a hydrophobic residue in the latch loop in HPGM1 led to a reduced enzyme activity (3.3% of the control) and caused disease in heterozygote patients.^39^ These findings are in line with our physiological characterization of the Δ*pgm1*+PGM1S47A strain and suggest that the regulatory mechanism discovered in this study is conserved in PGM1 enzymes.

This work contributes to the understanding of the regulation of PGM1 activity, which plays a pivotal role in the central carbohydrate metabolism. We showed the existence of a phosphorylation event that modulates PGM1 activity to control glycogen synthesis and degradation, a regulatory mechanism that seems to be essential for survival of nutrient deprivation and evolutionary conserved from bacteria to humans. The molecular mechanisms that lead to this post-translational modification remain, nevertheless, to be deciphered.

## Methods

### Cyanobacterial cultivation

The cyanobacterial strains used in this study are listed in **Table S1**. All strains were grown in BG_11_ supplemented with 5 mM NaHCO_3_ for vegetative growth, as described previously.^40^ Nitrogen starvation was induced as previously described by a 2-step wash with BG_11_-0 medium supplemented with 5 mM NaHCO_3_, which contains all BG_11_ components except for NaNO_3_.^7,41^ Resuscitation was induced by addition of 17 mM NaNO_3_ to cells residing in BG_11_-0. Cultivation was performed with continuous illumination (50 to 60 μmol photons m-2 s-1) and shaking (130 to 140 rpm) at 27 °C. Mutant strains were cultivated with the appropriate concentration of antibiotics.^8^ All strains used for this study are shown in **Table S1**. Biological replicates were inoculated from the same pre-cultures, but propagated, nitrogen-starved and resuscitated independently in different flasks under identical conditions.

### Protein overexpression and purification

*Escherichia coli* Rosetta-gami (DE3) was used for the overexpression of all proteins. All primers and plasmids used for protein overexpression are shown in **Table S2** and **Table S3**, respectively. All enzymes were Strep-tagged at the C-terminus. Cells were cultivated in 2xYT (1L of culture in 5L flasks) at 37 °C until they reached exponential growth (OD_600_ 0.6-0.8) and protein overexpression was then induced by adding 75 μg/L anhydrotetracycline, followed by incubation at 20°C for 16 h. Cells were harvested by centrifugation at 4000 g for 10 min at 4 °C, and disrupted by sonication in 40 mL of lysis buffer (100 mM Tris-HCl pH 7.5, 150 mm KCl, 5 mM MgCl_2_, DNAse, and cOmplete™ protease inhibitor cocktail (Roche, Basel)). The cell lysates were centrifuged at 20,000 g for 1 h at 4°C and the supernatants were filtered with a 0.22 μM filter.

The cell extracts were loaded onto a five-milliliter Ni-NTA Strep-tactin^®^ superflow column (Qiagen, Maryland, USA), washed with wash buffer (100 mM Tris-HCl pH 7.5 and 150 mm KCl), and eluted with elution buffer (100 mM Tris-HCl pH 7.5, 150 mm KCl, and 2.5 mM desthiobiotin). The buffer of all purified proteins was exchanged via dialysis using dialysis buffer (100 mM Tris-HCl pH 7.5, 150 mm KCl, and 5 mM MgCl_2_) and a 3 kDa cutoff dialysis tube. All purifications were checked via SDS-PAGE.

### Measurement of PGM1 activity in cell extracts

To determine the PGM1 activity in *Synechocystis* cell extracts an assay was adapted from Osanai et al.^42^ The PGM1 reaction was coupled to the G6PDH reaction and the glucose 6-phosphate-dependent conversion of NADP^+^ to NADPH was monitored by measuring the absorbance at 340 nm. Cells were harvested by centrifugation at 4000 g for 10 min at 4 °C, resuspended in lysis buffer (100 mM Tris-HCl pH 7.5, 10 mM MgCl_2_) and disrupted by using a “FastPrep®-24”(MP Biomedicals). The lysate was centrifuged for 10 min at 4 °C before the protein content was determined. When indicated. Approximately 50 μg of protein were used for each reaction. The reaction buffer was composed of 100 mM Tris-HCl pH 7.5, 10 mM MgCl_2_, 1 mM NADP^+^, 1 mM DTT and 1 U/mL G6PDH from *Saccharomyces cerevisiae* (G6378, Sigma Aldrich, Missouri, USA). The reaction was started by the addition of 10 mM glucose-1P. Absorption change at 340 nm was continuously measured for 15 min at 30 °C. As a blank, the change in absorption in the absence of glucose-1P was also measured and subtracted from the experimental values. The enzymatic activity was then calculated. At least three biological replicates were measured.

### Measurement of PGM1 activity *in vitro*

The reaction buffer was composed of 100 mM Tris-HCl pH 7.5, 150 mM KCl, 10 mM MgCl_2_, 1 mM DTT, 1 mM NADP^+^, 40 μM glucose-1,6-bisphosphate, and 1 U/mL G6PDH from *Saccharomyces cerevisiae* (G6378, Sigma Aldrich). 100 ng of C-terminus Strep-tagged PGM1 were added to each reaction. The reaction was started by the addition of glucose-1P. Absorption change at 340 nm was continuously measured for 15 min at 30 °C. The enzymatic activity was then calculated. At least three replicates were measured.

### Glucose-phosphate quantification

4 mL of chlorotic and resuscitating (24 h after NaNO_3_ addition) cultures were collected (OD_750_ ~ 0.8). Cells were harvested by centrifugation at 18,000 g for 1 min at 4°C. Pellets were immediately frozen in liquid nitrogen. Cells were lysed by addition of 0.2 M HCl and incubation at 95 °C for 15 min. Lysates were centrifuged at 18,000 g for 10 min at room temperature, then the supernatants were transferred to clean 2 mL tubes. Samples were neutralized with 1 mL of 1 M Tris-HCl pH 8. A glucose-1P and glucose-6P calibration curve were prepared. NADP^+^, KCl, and MgCl_2_ were added to samples and standard solutions to a final concentration of 1 mM, 150 mM and 10 mM, respectively. The absorbance of samples and standards were measured at 340 nm (blank measurement). 3 U of G6PDH from *Saccharomyces cerevisiae* (G6378, Sigma Aldrich) were added to all samples and standards and their absorbance at 340 nm was measured after incubation for 5 min at room temperature (glucose-6P measurement). 3 U of Pgm from rabbit muscle (P3397, Sigma) were added to all samples and glucose-1P standards and their absorbance at 340 nm was measured after incubation for 5 min at room temperature (glucose-1P measurement). The blank measurements were subtracted from the glucose-6P measurements, and the glucose-6P standard curve was used to determine the concentration of glucose-6P in the samples. The glucose-6P measurements were subtracted from the glucose-1P measurements, and the glucose-1P standard curve was used to determine the concentration of glucose-1P in the samples. Data were normalized to the OD_750_ of the sampled cultures. Three biological replicates were measured.

### Clear native PAGE – Immunoblot analysis

Proteins from 20 mL cell culture aliquots were lysed in extraction buffer with or without phosphatase inhibitors (50 mM Tris/HCl, 5 mM EDTA, 5 mM sodium fluoride, 5 mM sodium orthovanadate, pH 7.4) with a RiboLyser, conducting five cycles with a speed of 6.5 m s-1 for 15 s at 4°C. When indicated, cell extracts were treated with 1 U/mL alkaline phosphatase for 10 min at room temperature. Protein concentration was measured by Bradford assay and 10 μg total protein content was separated by non-denaturing clear native polyacrylamide gel electrophoresis (PAGE). Samples were mixed with loading buffer, loaded onto tris-glycine 8% separating native polyacrylamide gels, and run at 30 mA until the die reached the bottom of the gel. Protein transfer was performed on a semi-dry blotting system (peqlab) onto polyvinylidene fluoride (BioTrace™ PVDF, Pall Corporation) membranes at 20 V for 30 min. Membranes were blocked overnight in TBS-T buffer (50 mM Tris, 150 mM NaCl, 0.1 % Tween-20, pH7.5) and 5% milk powder at 4°C. Membranes were washed with TBS-T buffer and incubated with polyclonal primary antibodies against PGM1 (serum, produced in rabbit) for 1 h at RT in a 1:5000 dilution in TBS-T buffer. Membranes were washed again and incubated with peroxidase coupled secondary antibodies (Sigma A6154) for 1 h at RT in a 1:2000 dilution in TBS-T buffer. Proteins were visualized by Lumi-Light Plus detection reagent (Roche).

### Phylogenetic analysis

Sequences of diverse members of the α-D-PHM superfamily were obtained from NCBI RefSeq_protein database. Alignment and phylogenetic tree data were obtained using NCBI Constraint-based Multiple Alignment Tool (COBALT).^43^ Phylogenetic tree was generated with the Interactive Tree Of Life (iTOL) v5 online tool.^44^

### Recovery assay

Serial dilutions of chlorotic cultures were prepared (10^0^, 10^-1^, 10^-2^, 10^-3^, 10^-4^ and 10^-5^) starting with an OD_750_ of 1. 5 μl of these dilutions were dropped on solid BG_11_ agar plates and cultivated at 50 μmol photons m^-2^ s^-1^ and 27 °C for five days.

### Glycogen determination

Glycogen content was determined as described by Gründel et al.^29^ with modifications established by Klotz et al.^7^ Two-milliliter samples were collected, span down and washed with distilled water. Cells were lysed by incubation in 30% KOH at 95°C for 2h. Glycogen was precipitated by addition of cold ethanol to a final concentration of 70% followed by an overnight incubation at −20 °C. The precipitated glycogen was pelleted by centrifugation at 15000 g for 10 min and washed with 70% ethanol and 98% absolute ethanol, consecutively. The precipitated glycogen was dried and digested with 35 U of amyloglucosidase (10115, Sigma Aldrich) in 1 mL of 100 sodium acetate pH 4.5 for 2 h. 200 μl of the samples were mixed with 1 mL of 6% O-toluidine in acetic acid and incubated at 100 °C for 10 min. Absorbance was then read at 635 nm. A glucose calibration curve was used to determine the amount of glycogen in the samples. For every condition, at least three biological replicates were measured.

### Statistical analysis

Statistical details for each experiment can be found in the figure legends. Samples taken from cultures that were inoculated with the same pre-cultures, but propagated, nitrogen-starved and resuscitated independently in different flasks under identical conditions were considered different biological replicates. GraphPad PRISM was used to perform one-way ANOVA to determine the statistical significance. Asterisks in the figures were used to symbolize the p-value: One asterisk represents p ≤ 0.05, two asterisks p ≤ 0.01, three asterisks p ≤ 0.001, and four asterisks p ≤ 0.0001.

## Abbreviations

PGM1: phosphoglucomutase 1
α-D-PHM: α-D-phosphohexomutase
PMM: phosphomannomutase
PGM/PMM: phosphogluco/phosphomanno mutase
PNGM: phosphoglucosamine mutase
PAGM: phosphoacetylglucosamine mutase
PGM2/PGM2L1: phosphopentomutase/glucose-1,6-bisphosphate synthase

## Author contributions

S.D. performed cultivation experiments, cloning, protein purification, enzymatic assays, phylogenetic analysis, glycogen and glucose-phosphate determinations, immunoblot analysis, constructed mutants, analyzed data, and wrote manuscript with input from all authors. N.N. contributed cloning and sequence analysis. K.F. conceived study, interpreted data and edited manuscript.

## Acknowledgements

We thank Dr. Libera Lo Presti for her assistance writing this manuscript. This work was supported by the German Research Council (DFG) FOR 2816 “The Autotrophy-Heterotrophy Switch in Cyanobacteria: Coherent Decision-Making at Multiple Regulatory Layers”. Additionally, we acknowledge the infrastructural support from EXC 2124 “Controlling Microbes to Fight Infections” (ID 390838134).

## Data availability

Enzymatic data generated during this study have been deposited on the data and model management platform FAIRDOMHub^45^ and will be made publicly available upon publication of this work.

## Notes

**Conflicts of interest:** The authors declare no conflicts of interest.

### Competing Interest Statement

The authors have declared no competing interest.

### Summary of Updates

The manuscript has been revised after the identification of an annotation error of the PGM1 sequence of Synechocystis sp. PCC 6803 in Cyanobase. The new version focuses on the regulation of PGM1 activity via phosphorylation at a peripheral residue.

