## Supplementary information for "Phosphorylation of phosphoglucomutase 1 on a peripheral site tunes its activity to regulate glycogen metabolism"

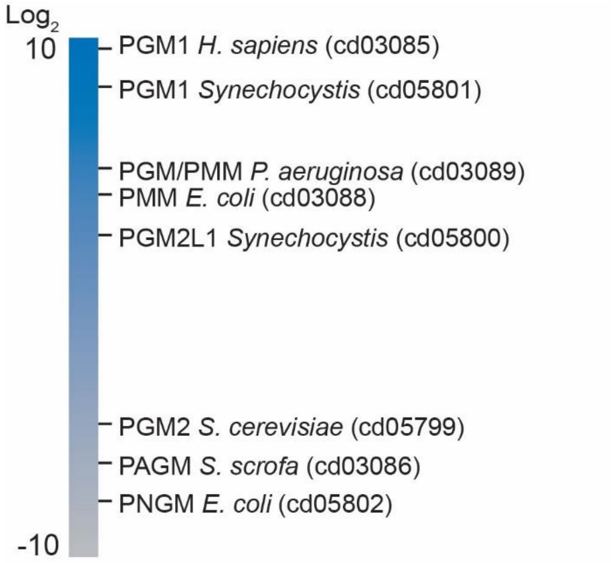


**Figure S1. Phosphoglucomutase activity of representative members of the different α-D-PHM subfamilies.** Related to Figure 3. Specific activity (U/mg) of different PHM enzymes using glucose-1-phosphate as a substrate was collected from the literature and is shown as $log_2$ and encoded in a color code from grey ($2^{-10}$) to blue ($2^{10}$).[11,22,23,24,25,26,27]

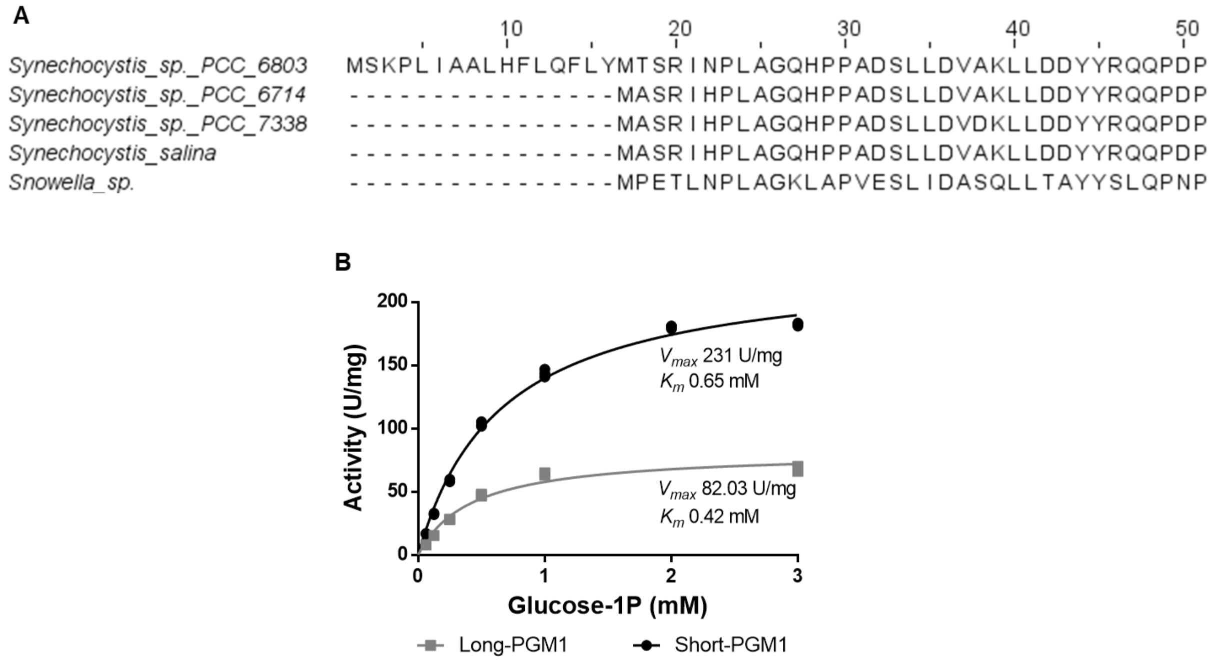


**Figure S2. Incorrect annotation of PGM1.** Related to Figure 3. (A) Alignment of PGM1 sequences from different *Synechocystis* strains. (B) Enzymatic characterization of the two versions of PGM1. Long-PGM1 and Short-PGM1 represent the enzyme the enzyme from *Synechocystis* sp. PCC 6803 with or without the 16-amino acid N-terminal extension that is absent in the annotation of the other strains, respectively.

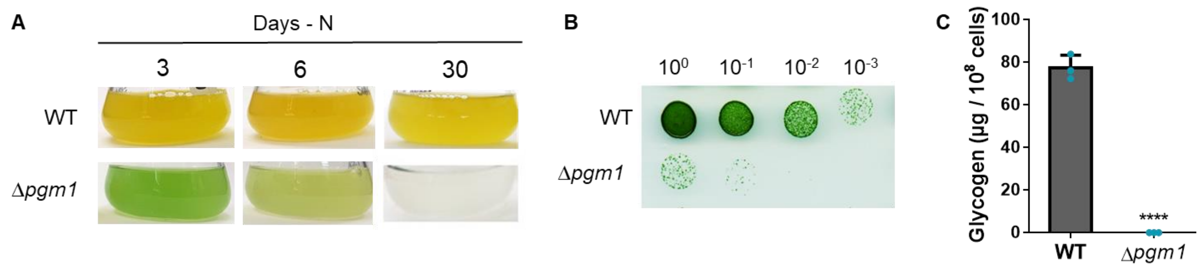


**Figure S3. Characterization of a PGM1 knockout strain (Δpgm1).** Related to Figure 4. (A) Pictures of WT, and Δ*pgm1* cultures after 3, 6, and 30 days of nitrogen starvation. (B) Recovery assay on a $BG_{11}$-agar plate of WT and Δ*pgm1* after 7 days of nitrogen starvation. Numbers on top represent the dilution factor, starting with an $OD_{750}$ of 1. Pictures were taken 5 days after dropping chlorotic cells on the plate. (C) Glycogen content of WT and Δ*pgm1* after 7 days of nitrogen starvation. Three biological replicates were measured. Error bars represent the SD; asterisks represent the statistical significance.

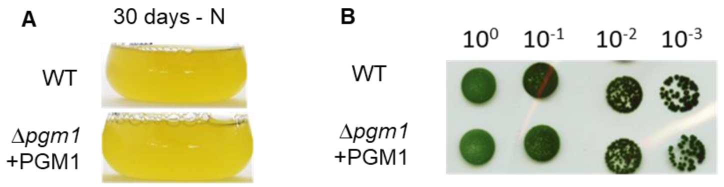


**Figure S4. Complementation of Δ*pgm1* with PGM1.** Related to Figure 4. (A) Pictures of WT and Δ*pgm1*+PGM1 cultures after 30 days of nitrogen starvation. (B) Recovery assay on a $BG_{11}$-agar plate of WT and Δ*pgm1*+PGM1. Numbers on top represent the dilution factor, starting with an $OD_{750}$ of 1. Pictures were taken 8 days after dropping chlorotic cells on the plate.

**Table S1. List of used strains**

| **Strain** | **Genotype** | **Purpose** |
|---|---|---|
| *E. coli* NEB10β | Δ(ara-leu) 7697 araD139 fhuAΔlacX74 galK16 galE15 e14-Φ80dlacZΔM15 recA1 relA1 endA1 nupGrpsL(StrR) rphspoT1 Δ(mrr-hsdRMS-mcrBC) | Molecular cloning |
| *E. coli Rosetta gami (DE3)* | Δ(ara-leu)7697 ΔlacX74 ΔphoA PvuII phoR araD139 ahpC galE galK rpsL (DE3) F'[lac$^+$ lacI$^q$ pro] gor522::Tn10 trxB pLysSRARE (Cam$^R$, Str$^R$, Tet$^R$) | Protein expression |
| Synechocystis sp. PCC 6803 GT | Wild-type | Control strain |
| Δ*pgm* | *Sll0726::Spec$^R$* | Characterization |
| Δ*pgm* + Pgm1 | Δ*pgm pVZ322(sll0726) Kan$^R$,Gen$^R$* | Characterization |
| Δ*pgm* + Pgm1S47A | Δ*pgm pVZ322(sll0726(S47A)) Kan$^R$,Gen$^R$* | Characterization |
| WT+Pgm1S47A | *pVZ322(sll0726(S47A)) Kan$^R$,Gen$^R$* | Characterization |

**Table S2. List of used primers**

| Primer | Sequence (5’ - 3’) |
|---|---|
| pASKC Lpgm1 fw | TAGAAATAATTTTGTTTAACTTTAAGAAGGAGATATACAAGTGTCTAAGCCCCTGATCGC |
| pASKC Lpgm1 rev | GTCTTATTTTTCGAACTGCGGGTGGCTCCAGCTAGCCATGCCCAAAGCCGAGGTAACAATG |
| pASKC Spgm1 fw | TAGAAATAATTTTGTTTAACTTTAAGAAGGAGATATACAAATGACAAGCAGAATTAATCC |
| pASKC Spgm1 rev | TCTTATTTTTCGAACTGCGGGTGGCTCCAGCTAGCCATGCCCAAAGCCGAGGTAACAATG |
| Pgm1 S47A fw | CTTTGGTACCgctGGCCATCGGG |
| Pgm1 S47A rev | CTCACTAACTGGGCGGGATTTTC |
| Pgm1 S47D fw | CTTTGGTACCgatGGCCATCGGGG |
| Pgm1 S47D rev | CTCACTAACTGGGCGGGA |
| pUC19-DSpgm1 fw | CCATGATTACGCCAAGCTTGCATGCCTGCAGGTCGACTGGCACAGCCATAGCCTAAAG |
| SmR-DSpgm1 rev | GTTTCTACAAACTCCATATGTCGACCCGGTCAATGTTCCG |
| DSpgm1-SmR fw | CGGAACATTGACCGGGTCGACATATGGAGTTTGTAGAAACG |
| USpgm1 SmR rev | CCTTTTTGGCTAGGATTTGGGAGCTCTTGACCGAACGCAG |
| SmR-USpgm1 fw | CTGCGTTCGGTCAAGAGCTCCCAAATCCTAGCCAAAAAGG |
| pUC19-USpgm1 rev | AAACGACGGCCAGTGAATTCGAGCTCGGTACCCGGGGATCCAGGCAATCAATCCGCTCAG |
| pVZ322 pgm1 fw | GCTTCCAGATGTATGCTCTTCTGCTCCTGCAGGTCGACTTTAGCCCAAAGCCGAGGTAAC |
| pVZ322 pgm1 rev | TGAATGTTCCGTTGCGCTGCCCGGATTACAGATCCTCTAGCAGGCAATCAATCCGCTCAG |
| Hpgm S20D fw | ACCGGGTACCGATGGTCTGCGTAAAC |
| Hpgm S20D rev | TTCTGGTCCTGGTACGCC |

**Table S3. List of used plasmids**

| Plasmid | Purpose |
|---|---|
| pASK IBA5C-LongPgm1 | Expression of C-terminus Strep-tagged *Synechocystis* (long) PGM1 in *E. coli* |
| pASK IBA5C-ShortPgm1 | Expression of C-terminus Strep-tagged *Synechocystis* (short) PGM1 in *E. coli* |
| pASK IBA5C -pgm1S47A | Expression of C- terminus Strep-tagged *Synechocystis* PGM1 S47A in *E. coli* |
| pASK IBA5C -pgm1S47D | Expression of C- terminus Strep-tagged *Synechocystis* PGM1 S47D in *E. coli* |
| pASK IBA5C-HPgm1 | Expression of C- terminus Strep-tagged *H. sapiens* PGM1 in *E. coli* |
| pASK IBA5C-HPgm1S20D | Expression of C- terminus Strep-tagged *H. sapiens* PGM1 S20D in *E. coli* |
| pUC19-pgm1 | Replacement of *pgm1* for Spec$^{R}$ in *Synechocystis* sp. PCC 6803 |
| pVZ322-pgm1 | Complementation of *pgm1* deletion |
| pVZ322-pgm1S47A | Complementation of *pgm1* deletion and WT with PGM1 S47A mutant variant |
